# KLIPP - a precision CRISPR approach to target structural variant junctions in cancer

**DOI:** 10.1101/2023.05.10.540176

**Authors:** Huibin Yang, Radhika Suhas Hulbatte, Alan Kelleher, Natalie Gratsch, Yin Wang, Philip L. Palmbos, Mats Ljungman

## Abstract

Current cancer therapies typically give rise to dose-limiting normal tissue toxicity. We have developed KLIPP, a precision cancer approach that specifically kills cancer cells using CRISPR/Cas9 technology. The approach consists of guide RNAs that target cancer-specific structural variant junctions to nucleate two parts of a dCas9-conjugated endonuclease, Fok1, leading to its activation. We show that KLIPP causes induction of DNA double strand breaks (DSBs) at the targeted junctions and cell death. When cancer cells were grown orthotopically in mice, activation of Fok1 at only two junctions led to the disappearance of tumor cells in 7/11 mice. This therapeutic approach has high specificity for tumor cells and is independent of tumor-specific drivers. Individualized translation of KLIPP to patients would be transformative and lead to consistent and simplified cancer treatment decisions.

## Introduction

Chromosome rearrangements are common in cancers where they drive tumor formation and typically occur early in carcinogenesis. (*1-5*). These rearrangements are mostly clonal, meaning that if they are present in the primary tumor, they will be present in metastases arising from the primary tumor (*4, 5*). Recent whole genome sequencing (WGS) studies of metastatic cancers found that they harbor a median of ∼200 chromosomal structural alterations per cell across tumor types (*6, 7*). Despite being a well-known feature of cancer, discovered more than a hundred years ago (*8*), this common cancer hallmark has not yet been broadly exploited therapeutically in the clinic. Oncogene amplifications (*9, 10*) and oncogenic fusion genes (*11, 12*) have been targeted directly with Cas9 and single guide RNAs (sgRNAs) leading to reduced oncogene expression and tumor growth inhibition. However, since Cas9 endonucleases also are guided to DNA sequences in normal cells with subsequent DSB induction, these approaches are expected to damage normal tissues.

Structural variants consist of DNA sequences that have moved to new locations or been brought together due to deletions. The unique feature of structural variants is that DNA sequences that normally are separated from each other by some distance in normal cells are juxtaposed at structural variant junctions (SVJs) in tumor cells. Thus, if a sequence-specific targeting approach could be developed towards the DNA sequences flanking these junctions, the cancer-specific SVJs could be attacked while having limited or no harmful effects to normal cells. Here we present a new approach that we call KLIPP^1^, to specifically target SVJs in cancer cells by using a “split” endonuclease system consisting of Fok1-dCas9 (*13*) nucleated around SVJs through sequence-specific sgRNAs (Fig. 1a). When positioned next to each other, the Fok1 subunits can homodimerize which is required for its endonucleolytic activity leading to the induction of a double strand break (DSB). It has been shown that Fok1-dCas9 has high fidelity and low off-target activities *in vivo* (*14*).

**Figure 1.**
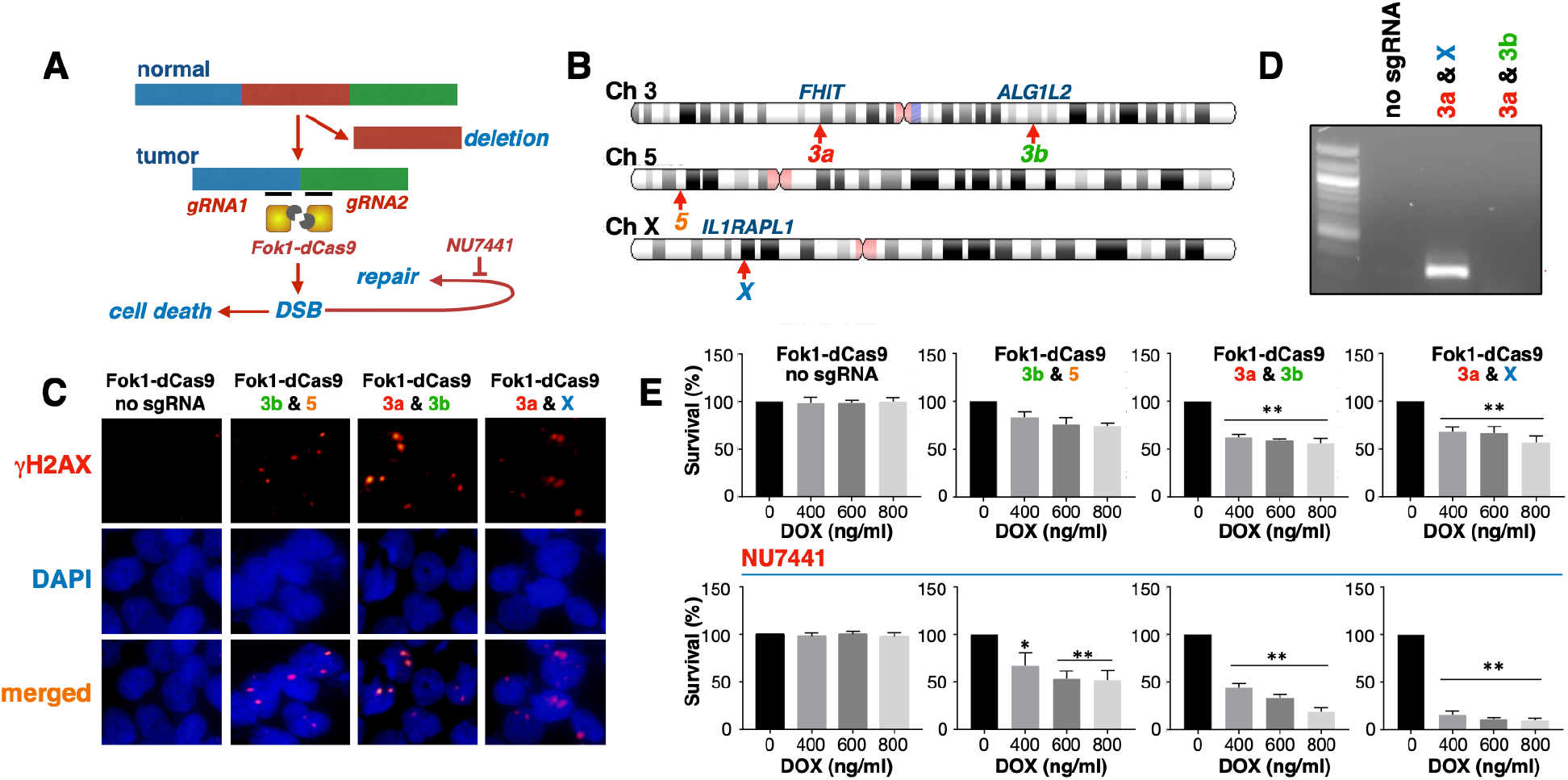
KLIPP induces DSBs and affects survival of HCT116 colon cancer cells. (**A**) Outline of the KLIPP approach targeting cancer-specific SVJs with sgRNAs and Fok1-dCas9 resulting in the dimerization and activation of the Fok1 endonuclease, the induction of DSBs, and cell death. Repair of DSBs can be suppressed by the DNA-PK inhibitor NU7441. (**B**) Chromosomal locations of the four SVJs selected from sequencing data from the HCT116 genomic sequence. Site 3a is a SVJ in the FHIT gene (active), site 3b is in the ALG1L2 gene (inactive), site 5 in a gene desert, and site X is in the IL1RAPL1 gene (active). (**C**) Induction of Fok1-dCas9 and SVJ-targeting sgRNAs by dox and inhibition of DNA-PK with 200 nM NU7441 for 24 h in HCT116 cells leads to the induction of DSBs as assessed using a γH2AX foci assay. (**D**) ChIP-PCR following immunoprecipitation using γH2AX-specific antibodies and primers specifically amplified sequences near the SVJ on chromosome X. (**E**) Induction of Fok1-dCas9 and SVJ-targeting sgRNAs causes reduced clonogenic survival that is augmented by the DNA-PK inhibitor NU7441. n=5, *p<0.05, **p<0.01.

We show that the KLIPP approach efficiently targets SVJ in both colon and bladder cancer cells in culture leading to induction of DSBs at sites of targeting and induction of cell death. No toxicity was observed when Fok1-dCas9 was expressed either alone or together with non-targeting sgRNAs. Using SVJ-targeting sgRNA pairs together with Fok1-dCas9 caused induction of DSBs, loss of clonogenic survival and induction of apoptosis. Assessment of the efficacy of KLIPP in an orthotropic mouse model of bladder cancer showed that the majority of cancers regressed and were undetectable at the end of the experiment. Thus, we show strong proof-of-concept that the KLIPP approach effectively targeted SVJs specifically in cancer cells both *in vitro* and *in vivo*. Successful development of this novel technology would be highly impactful and could be transformative in how we treat cancer

## Materials and Methods

### Cell culture

HCT116 cells were obtained from American Type Culture Collection and maintained in Dulbecco’s Modified Eagle Medium (Invitrogen) with 10% tet-free fetal bovine serum. Cells were grown in 100 mm dishes (Thermofisher) and incubated in a humidified chamber at 37°C with 5% CO_2_. UMUC3 cells were obtained from American Type Culture Collection and maintained in Dulbecco’s Modified Eagle Medium (Invitrogen) with 10% tet-free fetal bovine serum in 100 mm dishes (Thermofisher). Cells were incubated in a humidified chamber at 37°C with 5% CO2.

### Cloning of constructs

The plasmid vectors pX330A_Fok1-1x4 and pSLQ1685-dCas9-EGFP were purchased from Addgene. The dCas9 C-terminal portion-EGFP-PuroR cassette from the pSLQ1658-dCas9-EGFP vector was amplified by PCR and cloned into EcoR V/Sbf1 digested and gel purified pX330A_Fok1-1x4 vector by Gibson assembly. The engineered vector was sequence validated and named as pX330A_Fok1-dCas9-EGFP-2NLS-sgRNA. To create doxycycline-inducible Fok1-dCa9-GFP expression plasmid, TRE3G promoter from pLVX-TRE3G plasmid purchased from Clontech was PCR amplified with Kpn1 in forward primer and Aar1 in reverse primer. The PCR amplified TRE3G cassette was gel purified and cloned into Kpn1/Aar1 digested pX330A_Fok1-dCas9-EGFP-sgRNA plasmid replacing the CBh promoter to form pX330A_Fok1-dCas9-EGFP-sgRNA. To demonstrate inducible Fok1-dCas9-GFP expression, UMUC3 cells transfected with pX330A_Fok1-dCas9-EGFP plasmid together with pSV40_tet3G plasmid (a gift from Dr. Diane Simeone) were treated with different does 400, 600, and 800 ng/ml of doxycycline for 24 hours and Fok1-dCas9-GFP protein expression was validated both by western blot and immunofluorescent staining. G418 (Invitrogen) and 800 ug/µl of puromycin were used for selection of Fok1-dCas9 and sgRNA-expressing cells.

### Cell staining for χH2AX foci

HCT116 and UMUC3 cells were transferred to 8 well glass chamber slides and maintained in Dulbecco’s Modified Eagle Medium (Invitrogen) with 10% tet-free fetal bovine serum. Cells were treated with doxycycline (800 ng/ml) to induce the expression of Fok1-dCas9 and SVJ-targeting sgRNAs in the presence or absence of 200 nM NU7441 (Selleckchem). After 24 h at 37°C, the cells were fixed in a solution of 4% Paraformaldehyde in PBS and the slides were placed into each chamber for 30 minutes. Cells were washed with PBS 3 times and 100% methanol stored in -20 C° was added for 5 minutes and were then washed a final time. They were placed in 3% BSA-PBS-0.1% blocking buffer for 24 hours. The blocking buffer was aspirated and primary mouse γH2AX antibody (Invitrogen) was added to the blocking buffer in a ratio of 1:300 at 300 µl per well. The chambers were left at room temperature for 3 hours. These steps were then repeated using secondary goat-anti rabbit IgG Alexa Flur™ antibody (Invitrogen) in the same fashion. Slides were washed with PBS 5 times between each step. DAPI (Invitrogen) was applied for mounting and the coverslips were added. Imaging was performed using an Olympus U-HGLGPS fluorescent microscope with consistent exposure to assess expression of γH2AX and DAPI.

### γH2AX ChIP-PCR

We performed chromatin immunoprecipitation using cell lysates of doxycycline treated HCT116 and UMUC3 cells expressing dox-inducible Fok1-dCas9-sgRNA-GFP in the presence of 200 nM NU7441. After 72 h, cells were collected and the SimpleChIP plus Enzymatic Chromatin IP kit (Cell Signaling Technology, #9005) was used following the manufacturer’s protocol. The IP was performed using anti-γH2AX antibody (Abcam,ab2893). The PCR was executed with specific primers targeting sequences located ∼5 kb away from the CRJ of interest and each amplicon was 300-500 bp in length.

#### Primer sequence for HCT116-Xab

F primer: 5’CATTTCAGGCATATTAAGCACATT’3

R primer: 5’GGCAAGTCATATACCCAGAACAAC’3

#### Primer sequence for UMUC3-2ab

F primer: 5’GAGCAGGACAAGGTGCTTTC 3’

R primer: 5’GCACACTTCTTGGCATACGA 3’

### Clonogenic survival assay

HCT116 and UMUC3 cell lines with stable expression of doxycycline-inducible Fok1-dCas9-EGFP and two pairs of SVJ-targeting sgRNAs, were maintained in DMEM and RPMI1640 containing 10% tetracycline free FBS. Cells were trypsinized, washed two times with complete medium and ∼2000 cells were seeded on 60 mm Falcon dishes. After 24 h, cells were treated with doxycycline at concentrations of 0, 400, 600 and 800 ng/ml respectively in the presence or absence of 200 mM DNA-PK inhibitor NU7441. Cells expressing Fok1-dCas9 EGFP and non-targeting sgRNAs pairs served as control. Cells were maintained for 12 days and then fixed with 4% paraformaldehyde for 30 min and stained with crystal violet. Colonies containing at least 50 cells were counted as a colony and cell survival was determined as % of control cells.

### Assessment of apoptosis

UMUC3 cells expressing two pairs of SVJ-targeting sgRNA and Dox-inducible Fok1-dCas9-EGFP and UMUC3 cell-line expressing non-targeting sgRNA pairs (control) were grown and treated with 800 ng/ml of doxycycline for 72 h. The cells were collected and washed 2 times with PBS and fixed with ice cold 70% ethanol for 30 min at -20^0^ C. Cells were then stained with 1x PI (Sigma-Aldrich, #P-41070) after being washed 3 times with PBS. The samples were then immediately analyzed with flow cytometry (University of Michigan Flow Cytometry Core) and the fraction of cells in each sample containing sub-G_1_ DNA amounts was calculated.

### Lentiviral Transduction of UMUC-3 cells

UMUC-3 bladder cancer cells were transduced with Lenti6-GF1-CMV-VSVG lentivirus (vector Core, University of Michigan) for stable expression of luciferase. Briefly, cells were plated at 40-50% confluency in 6-well plates. 16-24 hours later, media was aspirated and 200 µl of 10X virus and 1 µl of polybrene were added with 2.8 mL DMEM (10% FBS) to cells. GFP expression was assessed under a fluorescent microscope after 72 hours. Each transduced cell line was analyzed every passage to ensure GFP expression remained 90-100% until orthotopic transplantation.

### In vivo establishment of orthotopic UMUC-3 tumors

Age matched NOD-scid IL2Rgamma^null^ (NSG) male mice were separated into two groups (20 per group). Luciferase labeled UMUC-3 bladder cancer cells (0.5 × 10^6^ cells/bladder), expressing two SVJ-targeting sgRNA pairs, were injected directly into the bladder of each NSG mouse as previously described (23, 24). Six days post-surgery, tumor formation was confirmed (T0) by bioluminescent imaging on IVIS Spectrum Optical Imaging System. Mice were separated into two groups where one group was administered 1 mg/ml doxycycline water (Sigma) (n=11) while the remaining mice continued to receive regular drinking water (n=10). Tumor growth was monitored weekly for all animals. The endpoint was predetermined at 28 days post-surgery (T3) at which point all mice were euthanized and bladders were recovered for endpoint weight measurement and imaging. Bladder tissue was formalin fixed, paraffin embedded and submitted for H&E staining (In vivo Animal Core, University of Michigan).

### Bioluminescence assessment of tumor growth

For weekly monitoring of tumor burden, mice were injected intraperitoneally with D-luciferin potassium salt at 120 mg/kg (Regis Technologies). Bioluminescent images were obtained using the IVIS Spectrum Optical Imaging System (Center for Molecular Imaging, University of Michigan) and analyzed with LivingImage^®^ software (Perkinelmer). For each timepoint, regions of interest (ROIs) were drawn around bioluminescent signal of each mouse and total flux measurements were automatically generated as integrated flux of photons (photons/sec).

### Staining of tumor tissues

For γH2AX staining, bladders bearing orthotopic UMUC3 tumors were harvested at 48, 72, or 96 hours post-doxycycline water administration and fixed in 10% NBF. Tissues were cryoprotected in a sucrose solution to prevent freezing artifact, OCT cryoembedded and sectioned. Paraffin-embedded mouse bladder tissues were sectioned (10 µm) and attached on glass slides. After Deparaffinization (xylene and ethanol) and rehydration, antigen unmasking was performed by boiling the slides submerged in 10 mM sodium citrate (pH 6.0) for 10 min in a pressured boiler. The slides were then blocked with PBS containing 5% BSA for 1 h in room temperature. Samples were incubated in primary antibody (anti-γH2AX, ab124781, Abcam) for 24 h at room temperature. After being washed thoroughly with PBS, the samples were incubated in secondary antibody (goat anti-rabbit IgG with Alexa Fluor 647, A21244, Invitrogen) for 1 h at 37°C. Both primary (1:100) and secondary (1:1000) antibodies were diluted in PBS containing 1% BSA. Hoechst 33342 (B2261, Sigma-Aldrich) was added into secondary antibody solution (120 µg/ml) to stain nuclei. After rinsing in deionized water, the slides were mounted with ProLong Diamond antifade mount (P36965, Invitrogen). Then cover glass was placed on slides and left in dark at room temperature for 24 h. Immunofluorescence pictures were taken by a confocal microscope (LSM800, Zeiss).

## Results

### Assessment of KLIPP targeting in HCT116 cells

We selected four SVJs from the Cosmic data collection (https://cosmic-blog.sanger.ac.uk) for the HCT116 colon cancer cell line that had favorable locations of PAM sequences, which are required for dCas9 positioning, and generated plasmids containing sequences for pairs of sgRNAs for each SVJ (Fig. 1B, Fig. S1). Combinations of two sgRNAs sequences were cloned into vectors expressing Fok1-dCas9-EGFP (Fig. S2). Due to the size limitation of these vectors, we were able to only fit 4 sgRNAs (two pairs) into each vector. Furthermore, we placed the Fok1-dCas9-EGFP fusion under the regulation of a tet-inducible promoter (TRE3G), which allows induction of expression by the addition of doxycycline (Dox) so that we could establish the cells in culture or as xenografts (Fig. S3). We also fused a EGFP domain to the Fok1-dCas9 protein so that we could better track its expression in cells (Fig. S4). The vectors were then used to transfect the HCT116 cells, generating sub lines for each tumor line stably expressing two SVJ-targeting sgRNA pairs and Dox-inducible Fok1-dCas9-EGFP (Fig. S5A). To assess whether induction of the Fok1-dCas9 alone or in combinations with SVJ-targeting sgRNA pairs resulted in the activation of the Fok1 endonuclease, we incubated the cells with Dox for 24 hours to induce Fok1-dCas9-EGFP and fixed and stained the cells for the presence of nuclear γH2AX foci which reports on the presence of a DSB. Expression of Fok1-dCas9 in the absence of any SVJ-targeting sgRNAs did not lead to any observable γH2AX foci (Fig. 1C). In contrast, cells expressing pairs of SVJ-targeting sgRNAs presented with one or two γH2AX foci, indicating successful targeting and Fok1 activation in many of the cells (Fig. 1C).

To verify that the DSBs were induced at the targeted SVJ, chromatin immunoprecipitation (ChIP) combined with PCR analysis with specific primers designed for sequences located near (∼5 kb) the targeted SVJs was performed. A strong amplification signal was found only in cells expressing the SVJ-targeting sgRNAs (Fig. 1D). We next explored cell fitness with a clonogenic assay using different concentrations of Dox that resulted in a dose-dependent induction of Fok1-dCas9-EGFP (Fig. S5B). Expressing Fok1-dCas9-EGFP alone did not affect clonogenic survival even at the highest dose of Dox used (Fig. 1E). In contrast, dose-dependent loss of clonogenic survival was observed for all three cell lines expressing SVJ-targeting sgRNAs and this effect was augmented when cells were incubated with the DNA-PK inhibitor Nu7441 (*15-17*) to inhibit DSB repair. (Fig. 1E). No effect of Nu7441 was observed when added to the control cells that expressed Fok1-dCas9-EGFP without targeting sgRNAs.

### Assessment of KLIPP targeting in UMUC-3 cells

To validate that KLIPP cell killing and specificity was not limited to HCT116 colon cancer cells, we performed similar experiments using the bladder cancer cell line UMUC-3. Genomic data for this cell line is available in the Cosmic data collection (https://cosmic-blog.sanger.ac.uk) we selected SVJs located on chromosome 2, 4 and 7 (Fig. 2A, Fig. S6) and generated UMUC-3 bladder cancer cells expressing sgRNA targeting pairs of these SVJs as well as dox-inducible Fok1-dCas9-EGFP. We also generated UMUC-3 cell lines expressing no or non-targeting sgRNAs. The non-targeting sgRNAs consisted of the two pairs of sgRNAs (3a & X) that effectively targeted SVJs present in HCT116 cells (Fig. 1), but since these SVJs are unique to HCT116 cells, these pairs of sgRNAs would not be expected to have any effect in the UMUC-3 cells. As expected, control cells expressing no sgRNA or non-targeting sgRNA formed no detectable γH2AX foci after inducing Fok1-dCas9 by Dox and incubating for 24 hours in the presence of NU7441 (Fig. 2B). In contrast, we observed many cells with two γH2AX foci per cell nucleus in cells expressing sgRNA targeting SVJs on chromosome 2 and 7, and one γH2AX focus per cells expressing sgRNA targeting SVJs on chromosome 4 and 7 (Fig. 2B). We conclude that the SVJs on chromosomes 2 and 7 were both successfully targeted by the selected sgRNAs while the selected SVJ on chromosome 4 did not respond. In concordance, induction of Fok1-dCas9-EGFP resulted in a stronger loss of clonogenic survival of the bladder cancer cells inducing two γH2AX foci (Ch. 2&7) compared to the cell line only inducing one γH2AX focus (Ch 4&7). Cell death was slightly increased in the cells expressing the SVJ-targeting sgRNAs when adding the DSB repair inhibitor NU7441 (Fig. 2C). Finally, we observed induction of apoptosis assessed by subG_1_ DNA content cells (Fig. 2D) and PARP1 cleavage only in cells expressing SVJ-targeting sgRNAs (Fig. S7).

**Figure 2.**
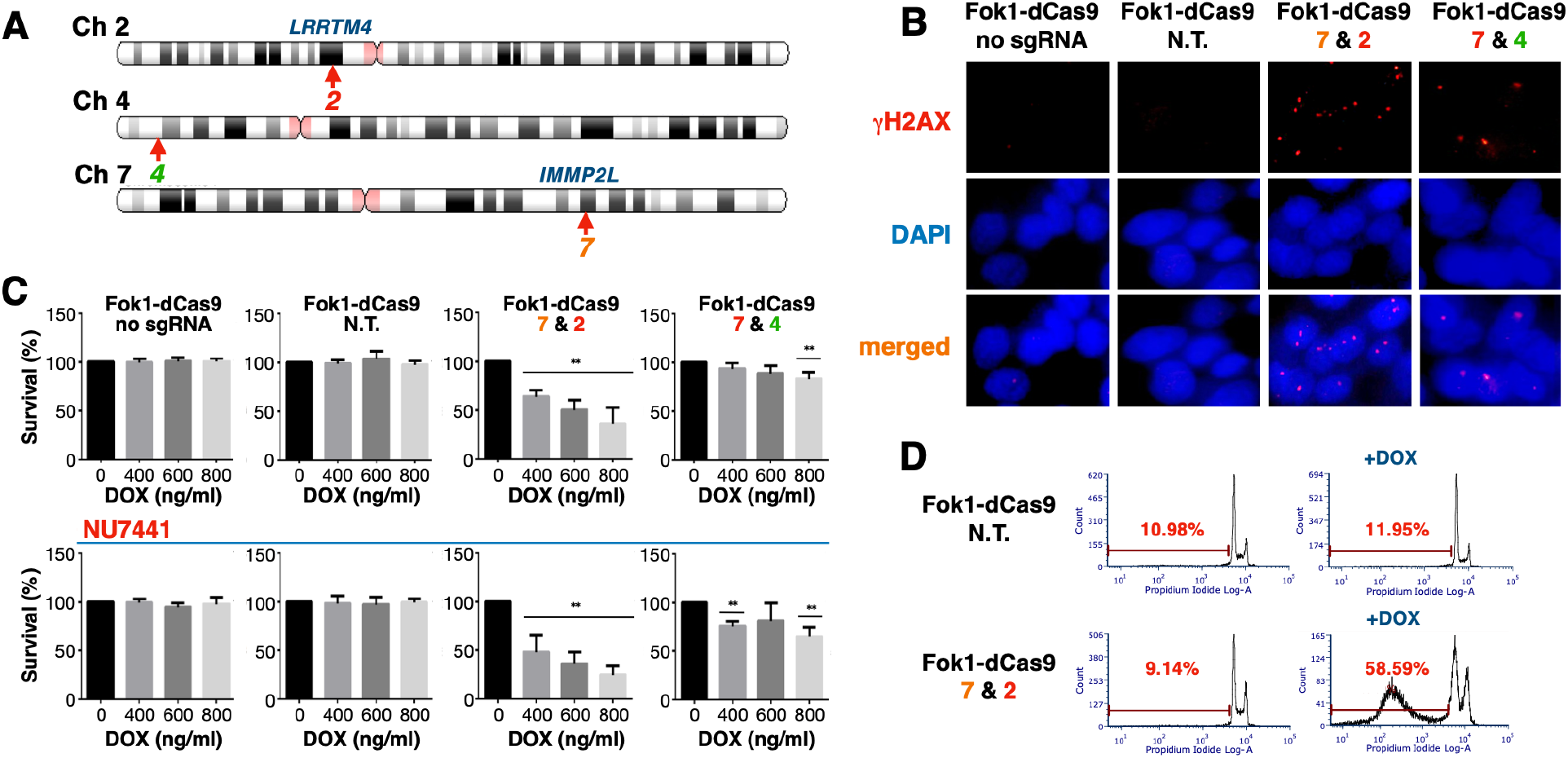
Induction of DSBs and toxicity by the precision CRISPR approach in UMUC-3 bladder cancer cells. (**A**) Chromosomal locations of the three SVJs selected from sequencing data from UMUC-3 cells. Number 2 is a SVJ in the gene LRRTM4 (inactive), number 4 is in a gene desert and number 7 is in the gene IMMP2L (active). (**B**) Induction of γH2AX foci following expression of SVJ-targeting sgRNA but not after expression of non-targeting sgRNAs after 24-hours of doxycycline treatment. (**C**) Different concentrations of doxycycline were used for 12 days in the absence (top) or presence (bottom) of the DSB repair inhibitor NU7441 and the numbers of colonies present was compared to control cells not treated with doxycycline. n=5, *p<0.05, **p<0.01. (**D**) Induction of apoptosis in UMUC-3 cells was assessed by subG_1_ DNA content determinations using propidium iodide staining and flow cytometry 72 hours after adding doxycycline to the media.

### Assessment of efficacy of KLIPP against orthotopic UMUC3 cells *in vivo*

We next grew the UMUC-3 cells expressing Dox-inducible Fok1-dCas9 in the presence or absence of SVJ-targeting sgRNA pairs as orthotopic xenografts in mice. These cells had been transfected with lentivirus carrying the luciferase gene so that the tumors that grew out could be monitored with bioluminescence imaging (Fig. 3A). Six days after implantation of the cancer cells, bioluminescence monitoring showed that all mice harbored a similar tumor burden (Fig. 3B). At this point, Dox was supplied in the drinking water of half of the mouse colony and the tumor burden was assessed with weekly bioluminescence imaging. Fixation and staining of tumors from two mice 96 hours after adding Dox to the drinking water provided evidence of induction of DSBs in the tumors as detected by γH2AX foci (Fig. 3D). Inspection of the bioluminescent signal of the tumors showed significantly reduced bioluminescence in the dox-treated group compared to mice not receiving Dox where bioluminescence increased dramatically over three weeks (Fig. 3B). When the bladders were extracted at the end of the experiment, the bladders from the Dox-treated animals were significantly smaller than the bladders in the control group suggesting a strong tumor growth inhibiting effect of inducing the Fok1-dCas9 and SVJ-targeting sgRNA pairs (Fig. 3F). Importantly, the Dox-treated animals did not present with gross metastases in the abdominal cavity, while the control animals presented with locally advanced bladder tumors, hemoperitoneum, and with multiple metastases (Fig. S8). Histologic examination demonstrated that while all the control bladders had extensive tumor burden, 7/11 (64%) of the Dox-treated animals had no detectable tumors (Fig. 3F). Thus, induced expression of Fok1-dCas9 and SVJ-targeting sgRNA pairs in orthotopic bladder tumor cells resulted in the induction of DSBs (γH2AX) and reduction or elimination of bladder tumors and metastases in vivo.

**Figure 3.**
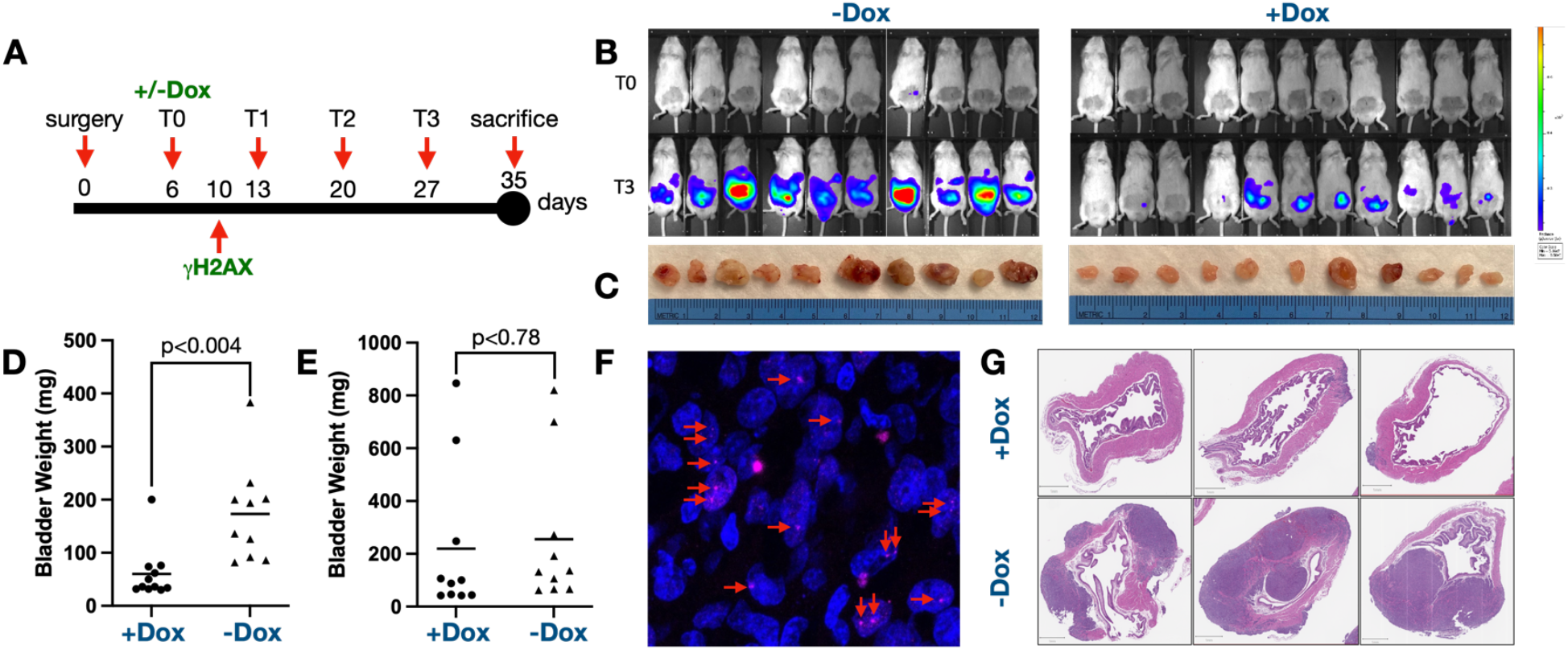
Induction of Fok1-dCas9 and SVJ-targeting sgRNAs in vivo suppresses tumor growth of orthotropic bladder cancers. (**A**) Experimental timeline of in vivo experiments. Doxycycline (Dox) was added to the drinking water of 13 mice at day 6 after implantation. Two of these mice were sacrificed 4 days later and tissue were stained for the presence of γH2AX. (**B**) Bioluminescence of orthotropic UMUC3 bladder cancer cells after 6 days of growth (T0) and after an additional 3 weeks (T3) without (left) and with Dox in the drinking water (right). (**C**) Bladders harvested at the conclusion of the experiment. During harvesting it was observed that mice not treated with Dox had many metastases in the abdominal cavity while Dox-treated animals did not. (**D**) Bladder weights of 11 mice carrying UMUC3 tumors with SVJ-targeting sgRNA (7&2) treated with Dox (left) and 10 mice not treated with Dox (right). The Dox-treated group had significantly lower average bladder weight than the untreated group. (**E**) Bladder weights of mice carrying UMUC3 tumors with Dox-inducible Fok1-dCas9 but without sgRNAs. Treatment with dox did not significantly affect the bladder weights of these mice. (**F**) Induction of γH2AX foci in vivo 4 days after adding Dox to the drinking water. (**G**) H&E staining of three bladders from the dox-treated animals (top) and three samples from non-dox treated animals (bottom) with UMUC-3 tumors with Dox-inducible expression of Fok1-dCas9 and two SVJ-targeting sgRNA pairs (7&2). The frequency of a complete response (CR), defined as no detectable bladder tumors, was 7/11 mice (64%) in the Dox-treated cells while all the non-treated animals had substantial tumors as did all the animals with UMUC3 tumors with Dox-inducible Fok1-dCas9 without sgRNAs.

## Discussion

While a few previous studies have used CRISPR technology to exploit gene amplifications and gene fusions in cancer cells, these approaches result in Cas9-induced cutting of genomes even in normal cells. The innovation of the KLIPP approach is that cancer cells are specifically targeted and killed by nucleating two parts of an endonuclease using pairs of sgRNAs designed to bind to both sides of SVJs unique to the cancer cells. Here we show that the Fok1-dCas9 approach led to efficient targeting of cancer cells both in cell culture and *in vivo*. Moreover, when sgRNA pairs that targeted SVJs in one cancer cell line were used in another, no DSBs or toxicity were observed. Due to tumor cell heterogeneity, it is likely that some of the SVJs selected for the treatment may not be present in all tumor cells and therefore cells lacking the targeted SVJs would be unaffected and could repopulate the tumor. Furthermore, cells may introduce mutations at the cleavage sites leading to the escape from further sgRNA binding and Fok1-dCas9 targeting. However, in contrast to drug resistance described for other targeted therapies, KLIPP can be applied iteratively using a new set of sgRNAs targeting SVJs present in an outgrown tumor. Additionally, WGS will identify the most prevalent SVJs across a clonally heterogeneous tumor that also should be the most effective SVJs to target.

The fact that as few as two DSBs induced by the Fok1 homodimers were sufficient to dramatically reduce tumor burden in our orthotopic bladder cancer model is especially encouraging. To expedite implementation of KLIPP as a therapy in the clinic, we are currently testing lipid nanoparticle formulations for the capturing and delivery of the KLIPP reagents to tumors in pre-clinical orthotopic cancer mouse models (*18-23*). The results presented here provide the proof-of-concept that the SVJ-targeting KLIPP approach can be used to treat and kill tumor cells *in vitro* and *in vivo*, a paradigm-shifting advance which would expedite the prospect of a universal and safe cancer treatment.

## Acknowledgements

We would like to acknowledge current and former members of the Ljungman lab for their contributions to this project, Dr. Analisa DiFeo for donating mice for these experiments, and Tom E. Wilson and Philip C. Hanawalt for reviewing this manuscript.

## Funding

This project has been supported by:

Private funds by Mr. Ron Weiser (ML),

Department of Radiation Oncology (ML)

Rogel Cancer Center (ML),

Research grants Mcubed, Kickstart and MTRAC from University of Michigan (ML)

National Cancer Institute grant CA213214 NCI (ML, PP)

Geri Fournier Ovarian Cancer Research Fund MIOCA (ML)

## Author contributions

Conceptualization: M.L.

Funding acquisition: M.L.

Investigation: H.Y., R.S.H., A.K., N.G., Y.W.\

Methodology: M.L., H.Y.

Project administration: M.L.

Supervision: M.L., P.P.

Visualization: M.L., P.P.

Writing original draft: M.L.

Writing, review and editing: M.L., H.Y., R.S.H, A.K., N.G. and P.P.

## Data and materials availability

All data are available in the main text and in the supplemental materials.

## Supplemental Materials

Supplementary Text

Figs. S1 to S8

## Footnotes

1 “KLIPP is a Swedish word that means “cut” and “opportunity”

