## Supplemental Figures for "KLIPP - a precision CRISPR approach to target structural variant junctions in cancer"

This PDF file includes:

Figs. S1 to S8

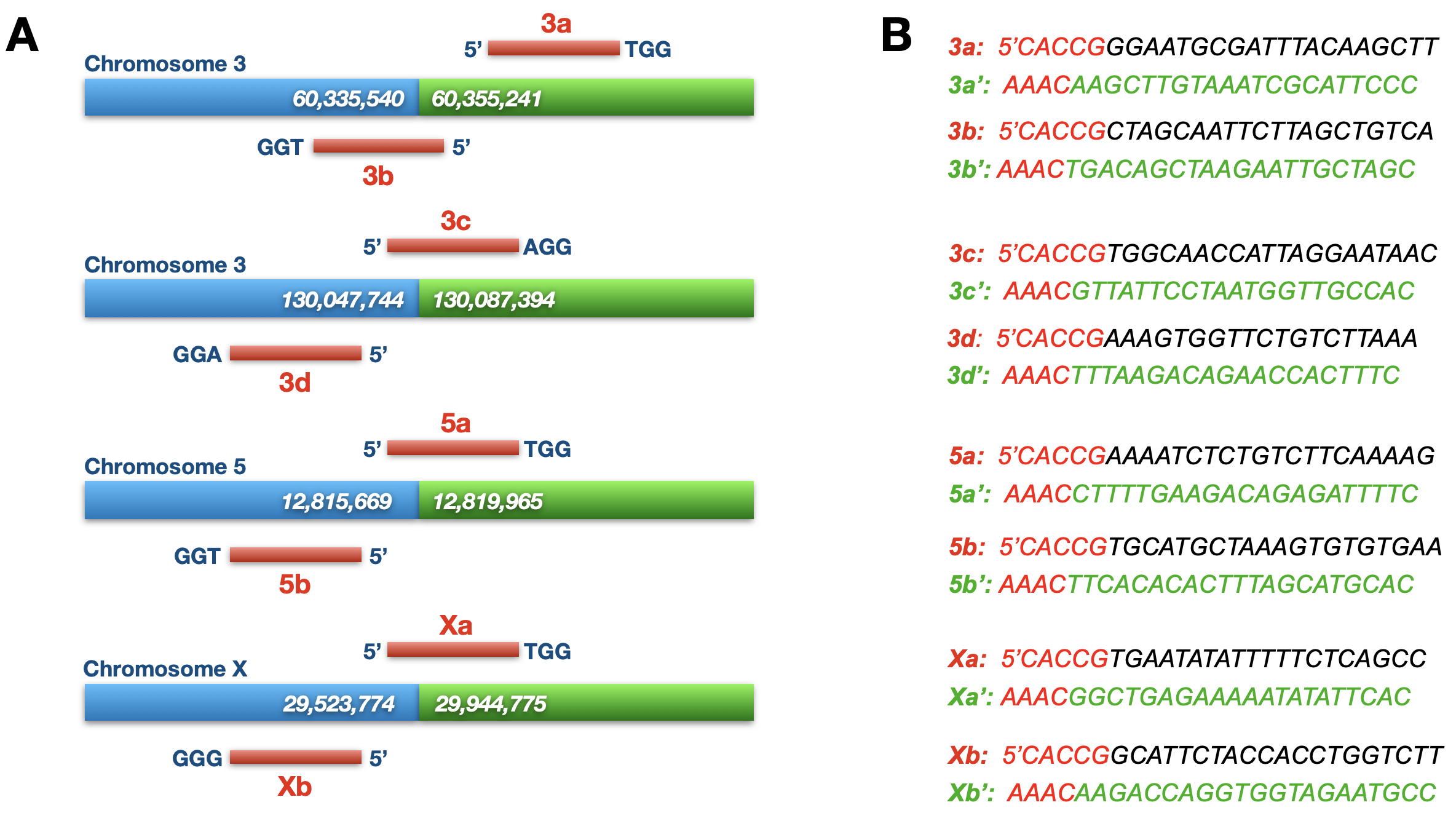

Fig. S1. (A) Location of the 4 SVJs in HCT116 cells selected for targeting. Coordinates for the breakpoints (COSMIC) are given as well as the approximate locations of the sgRNA pairs on either side of the CRJ. (B) Sequences of the sgRNAs as well as the reverse complementary sequences (‘) are shown.

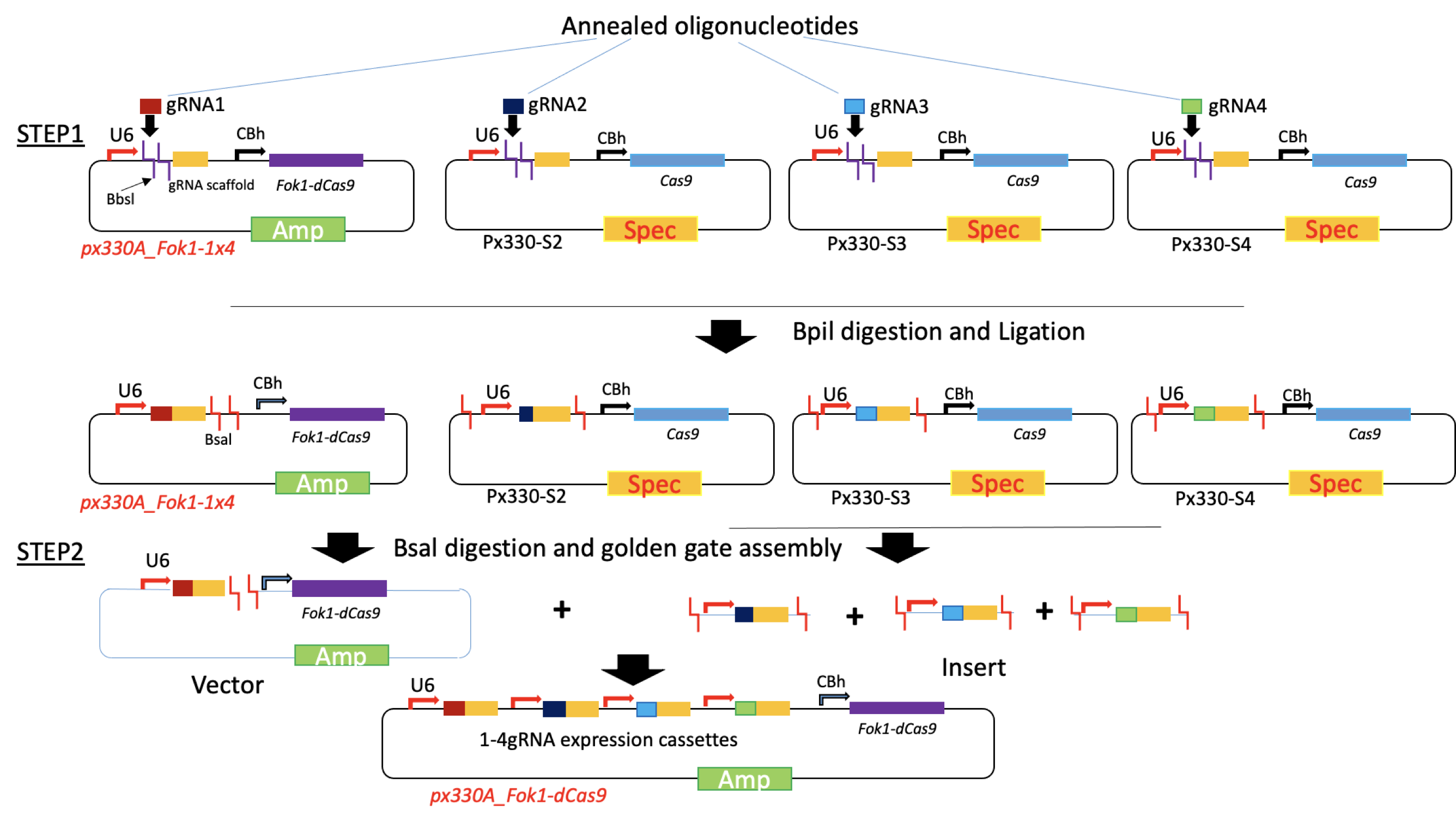

Fig. S2. Vectors of pX330A_Fok1-1x4, pX330S2, pX330S3 and pX330S4 purchased from Add gene were digested with Bpi1. The linearized vectors were gel purified. Then the annealed sgRNA 1 (5a, 3a for HCT116, 7a for UMUC-3) was cloned individually into pX330A_Fok1-1x4,  sgRNA2 (5b, 3b for HCT116, 7b for UMUC-3) into pX330S2, sgRNA3 (3c, Xa for HCT116, 2a, 4a for UMUC-3) into pX330S3, and sgRNA4 (3d, Xb for HCT116, 2b, 4b for UMUC-3) into pX330S4. Finally, sgRNAs (1-4) expression vectors were digested with Bsal and assembled into pX330A_Fok1-1x4 by golden gate assembly. The 4 sgRNAs assembled vectors were named as pX330A_FoK1 -5ab3cd, -3ab3cd, 3abXab (HCT116) respectively and pX330A_Fok1-7ab2ab and -7ab4ab (UMUC-3).

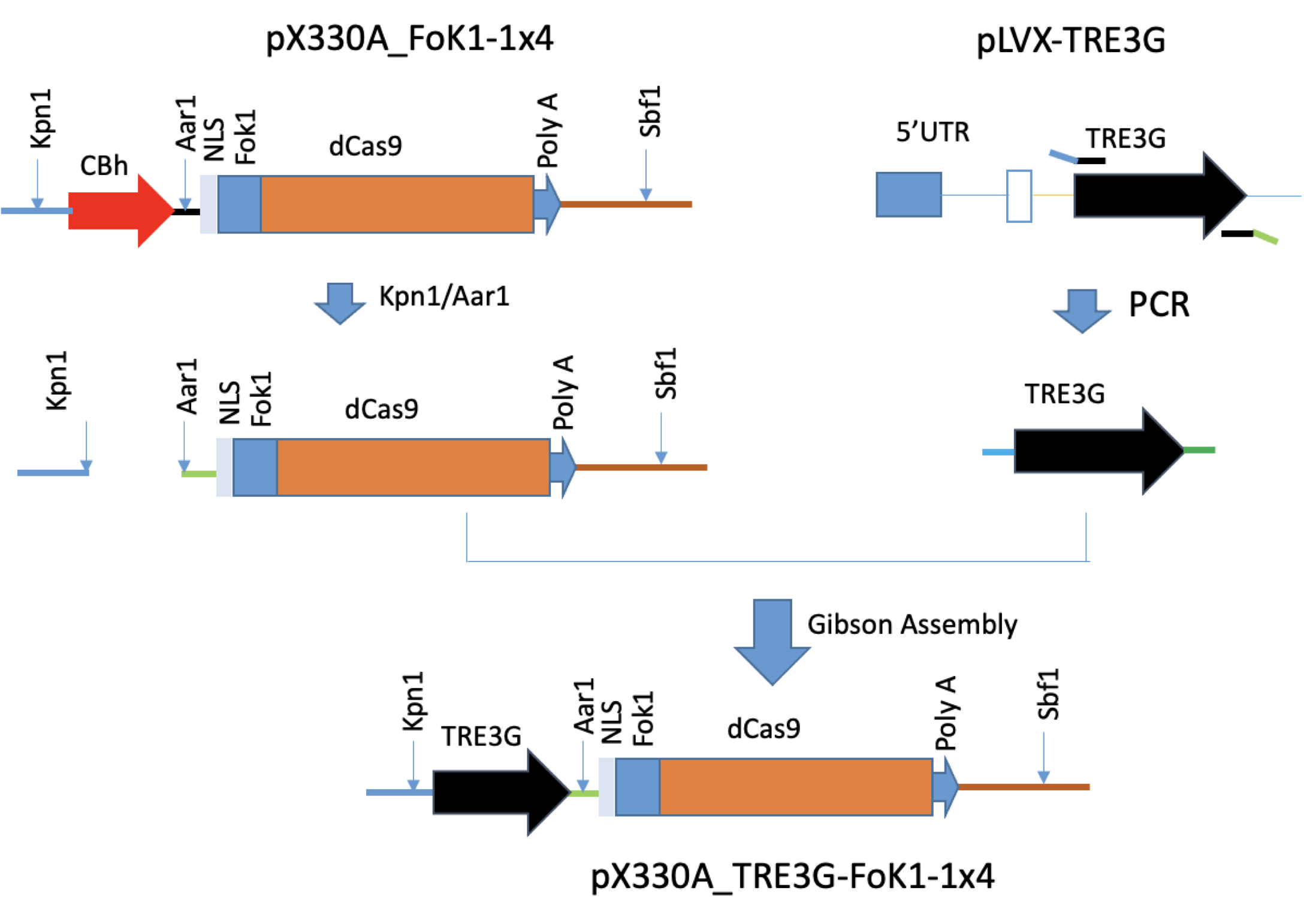

Fig S3. To create pX330_A-TRE3G-Fok1 plasmid, TRE3G from pLVX-TRE3G (purchased from Clontech) was PCR amplified with Kpn1 in forward primer and Aarl in reverse primer. The PCR amplified TRE3G cassette was cloned into Kpn1/Aarl digested pX330A_Fok1-sgRNAs plasmid replacing the CBh promoter to form pX330A_TRE3G-FoK1-sgRNA2 using Gibson assembly (New England Biolabs).

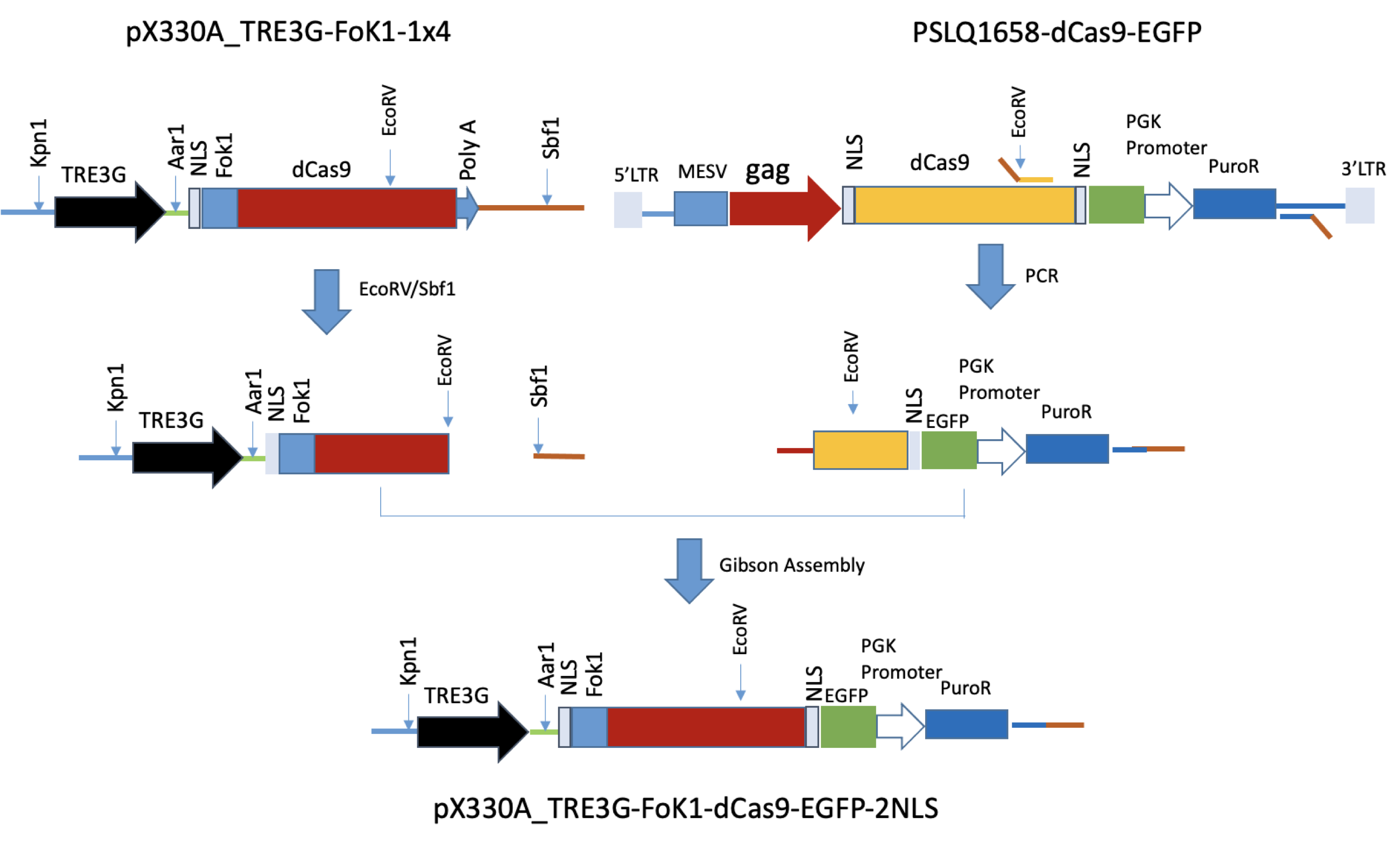

Fig. S4. To insert EGFP reporter at terminus of dCas9 of pX330A_Fok1-sgRNAs, dCas9 C-terminal part –NLS-EGFP-Puromycin resistant cassette was PCR-amplified and then cloned into EcoRV/Sbfl digested and gel purified pX330A_Fok1-sgRNAs vector by Gibson assembly (New England Biolabs). The engineered plasmid was named as pX330A_Fok1-dCas9-EGFP-2NLS-sgRNAs.

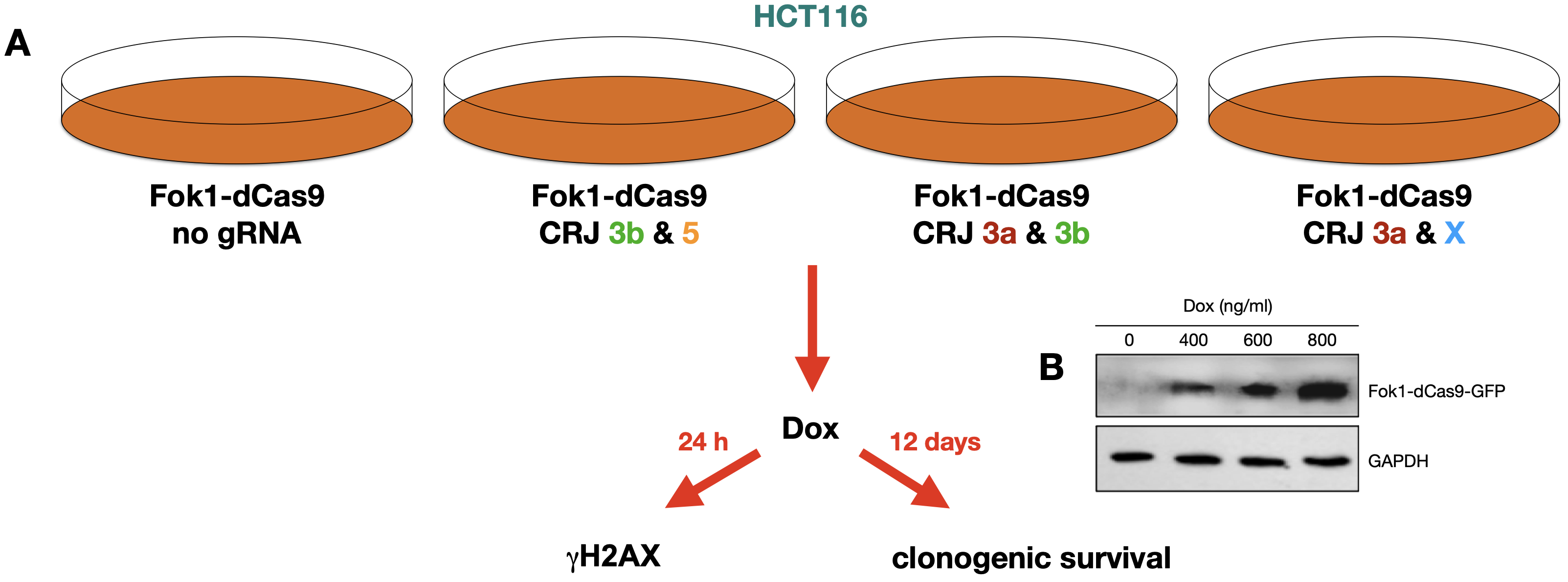

Fig. S5. (A) Experimental set up. HCT116 colon cancer cells were transfected with pX330A_Fok1-dCas9-EGFP-2NLS containing either no sgRNAs (left) or different combinations of pairs of sgRNAs targeting the SVJs shown in Fig. 1b. By adding doxycycline to the media, the Fok1-dCas9-EGFP cassette will be activated generating Fok1-dCas9 protein. Assessment of DSB induction is monitored by γH2AX foci 24 h after addition of doxycycline and the toxic effects are monitored using the clonogenic survival assay. (B) Cells were treated with different amount of Dox for 24 hours and the cell lysates were prepared for Western blot to measure Fok1-dCas9-EGFP expression using anti-GFP antibodies. GAPDH expression was used as a loading control.

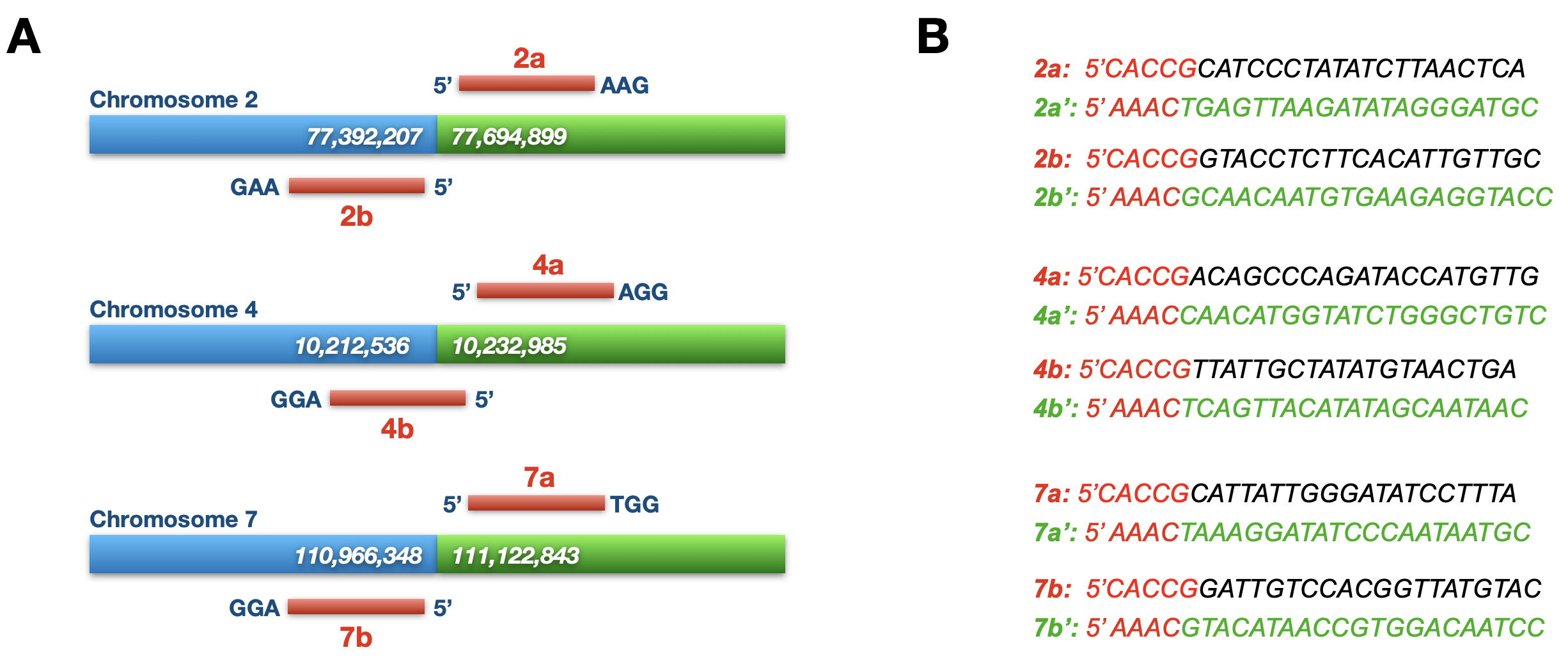

Fig. S6. (A) Location of the 4 SVJs in UMUC-3 cells selected for targeting. Coordinates for the breakpoints (COSMIC) are given as well as the approximate locations of the sgRNA pairs on either side of the SVJ. (B) Sequences of the sgRNAs as well as the reverse complementary sequences (‘) are shown.

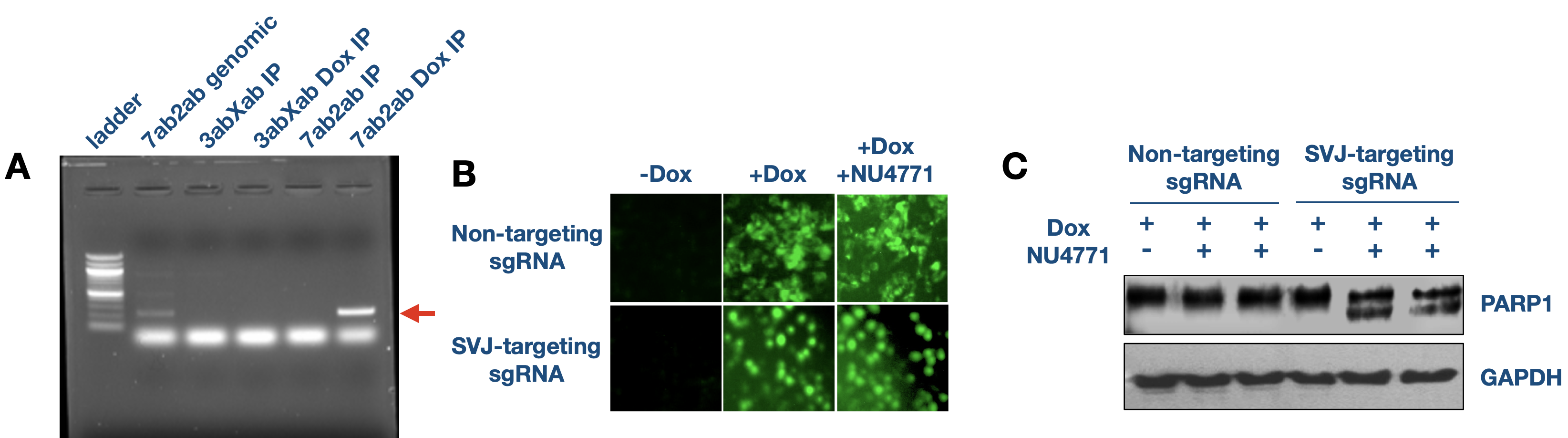

Fig. S7. (A) γH2AX-ChIP-PCR showing that Dox-induced expression of Fok1-dCas9 results in the specific induction of DSBs in UMUC-3 cells at sites of sgRNA binding (B) Addition of doxycycline induces a robust expression of Fok1-dCas9-EGFP within 24 hours. (C) Incubations of the cells with doxycycline for 72 hours results in PARP1 cleavage as an indication of apoptosis.

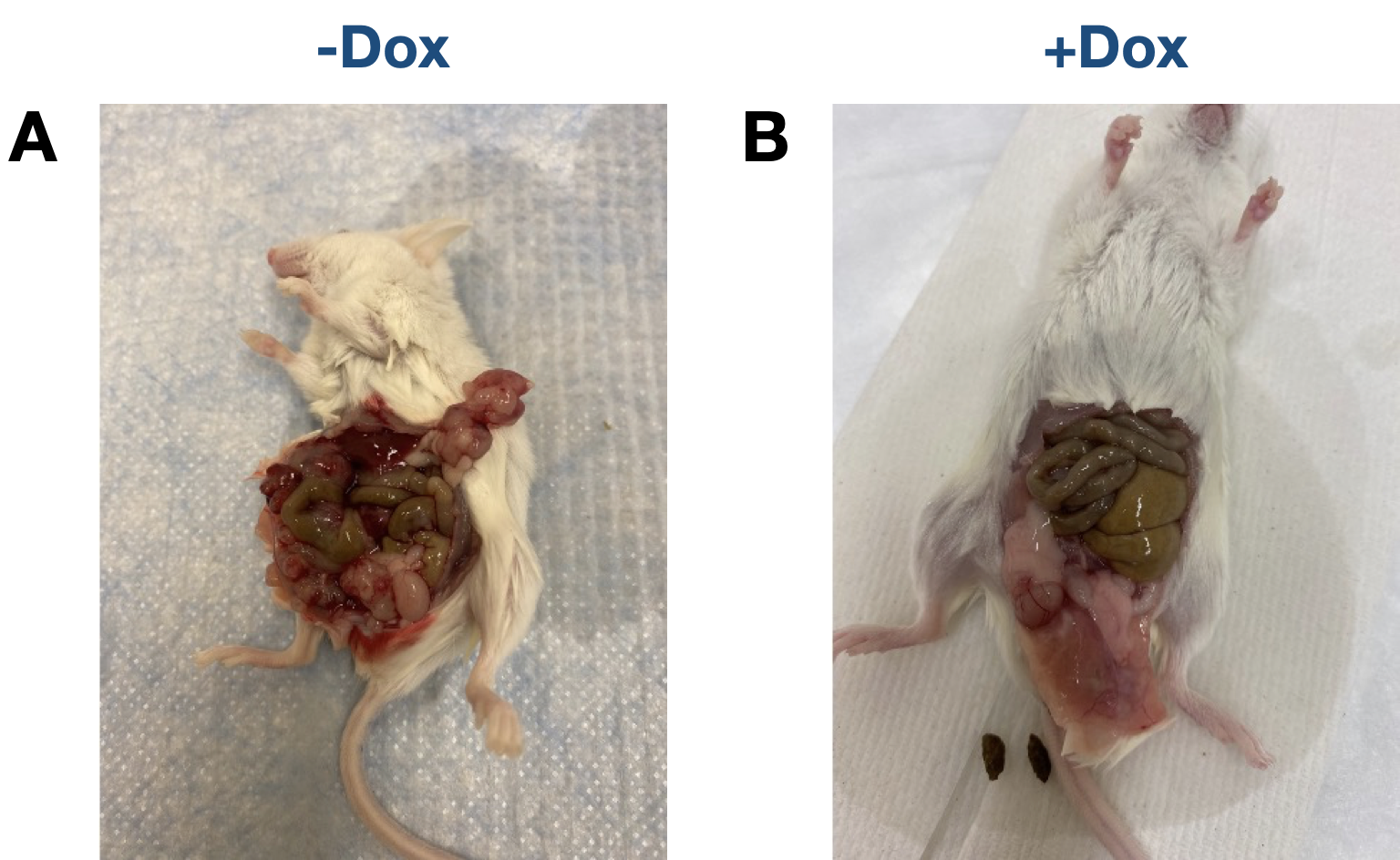

Fig. S8. (A) Mice not receiving Doxycycline in the water following implantation all showed evidence of metastasis in the abdomen at the termination of the experiment. (B) Mice receiving Doxycycline on day 6 after implantation and throughout the course of the experiment all showed normal abdomens without the presence of metastasis at the termination of the experiment.
